# Pericytes require physiological oxygen tension to maintain phenotypic fidelity

**DOI:** 10.1101/2024.08.28.606682

**Authors:** Tamara McErlain, Elizabeth C McCulla, Morgan J Glass, Lauren E Ziemer, Cristina M Branco, Meera Murgai

## Abstract

Pericytes function to maintain tissue homeostasis by regulating capillary blood flow and maintaining endothelial barrier function. Pericyte dysfunction is associated with various pathologies and has recently been found to aid cancer progression. Despite having critical functions in health and disease, pericytes remain an understudied population due to a lack of model systems which accurately reflect *in vivo* biology. In this study we developed a protocol to isolate and culture murine lung, brain, bone, and liver pericytes, that maintains their known phenotypes and functions. We demonstrate that pericytes, being inherently plastic, benefit from controlled oxygen tension culture conditions, aiding their expansion *ex vivo*. Primary pericytes grown in physioxia (10% O_2_ for lung; 5% O_2_ for brain, bone, and liver) also better retain pericyte phenotypes indicated by stable expression of characteristic transcriptional and protein markers. In functional tube formation assays, pericytes were observed to significantly associate with endothelial junctions. Importantly, we identified growth conditions that limit expression of the plasticity factor *Klf4* to prevent spontaneous phenotypic switching *in vitro*. Additionally, we were able to induce pathological pericyte phenotypic switching in response to metastatic stimuli to accurately recapitulate *in vivo* biology. Here, we present a robust method for studying pericyte biology in both physiology and disease.

## Introduction

Pericytes are critical components of microvascular homeostasis and are found across heterogeneous capillary beds^1,2^. Pericyte detachment or loss is associated with microvascular dysfunction observed in many pathologies, including aging, ischemic disease, neurodegeneration, and cancer^3^. Pericyte pathologies also extend beyond vascular dysfunction^4^, involving extracellular matrix (ECM) remodeling and fibrosis^5,6^. We and others have previously found that pericytes are critical to promoting cancer metastasis^7,8^.

Despite growing clinical interest in pericytes for their role in disease and regenerative medicine, the field has struggled to develop an *in vitro* toolkit for pericyte research that is reflective of physiology, relatively cost-effective, and high throughput^3^. This is in part due to the heterogeneity of origin^9,10^, function^2^, and molecular markers^1,11^, which has made it difficult not only to define pericytes but also to model them in a physiologically accurate way. The inherent phenotypic plasticity of pericytes, which allows them to differentiate into fat, muscle, cartilage, and bone cells^12–14^ may make them particularly sensitive to environmental cues. This sensitivity may in fact underly the inconsistences observed between pericyte isolation and culturing methods which in turn inevitably results in discrepancies in data obtained from different protocols, clouding reliability and translational value of the associated findings. Here, we sought to identify an *in vitro* pericyte modeling system that would most faithfully represent pericyte homeostatic and pathological phenotypes.

The optimal pericyte model should support expansion of cells for high throughput studies while accurately recapitulating pericyte biology without inducing spontaneous differentiation or pathological activation. The differentiation potential of pericytes presents a challenge for establishing reproducible cell culture models; however, similar difficulties have been overcome in the culture of pluripotent stem cells by regulation of oxygen tension^15–18^. *In vivo*, different tissues experience varying oxygen levels: lung tissue is exposed to oxygen partial pressures equivalent to 1-10% O_2_, while brain, bone, and liver tissues typically encounter 1-5% O_2_^19,20^. Standard tissue culture practice maintains cells at ∼21% O_2_ (hyperoxia); however, mimicking *in vivo* oxygen tension (physioxia) can more accurately reflect cell identity, function, and response to stimuli^19,20^. Physiological oxygen tension has been demonstrated to be an important factor of stem cell niches, functioning in cell fate determination with low oxygen tension being essential to maintain the plasticity and proliferation of stem cells^15–18^. Importantly it has been demonstrated that hyperoxia exposure impacts many cellular processes and limits the ability of *in vitro* tools to replicate *in vivo* biology^21,22^.

The functional role of pericytes as oxygen sensors^23,24^ together with their inherent plasticity, might suggest that this population is particularly sensitive to the oxygen tensions in which they are grown. Using the knowledge available in the microvascular field^25,26^, we developed a protocol for the isolation of murine pericytes for *in vitro* studies. We investigated if pericytes require physiological oxygen tension to maintain their homeostatic phenotype and response to pathological cues. Learning from the culturing practices of other pluripotent populations, in this study we sought to evaluate the understudied role of physiological oxygen tension to better model and retain pericyte phenotype and function. We report the development and characterization of a model that captures the phenotypic plasticity of pericytes observed in vascular pathologies, including cancer.

## Results

### Physiological oxygen levels support the growth of primary lung pericytes *ex vivo*

To examine pericyte responses to oxygen levels *in vitro*, NG2-expressing pericytes isolated from mouse lungs were cultured in hyperoxia (21% O_2_; standard tissue culture incubator) or physioxia (10% O_2_ for lungs). Phase contrast images were obtained at key time points during the expansion of pericytes, and proliferation was assessed at passage 1 **(Fig. 1A)**. To identify media conditions that support cell growth while maintaining pericyte biology, we developed a growth medium (GM) to promote pericyte culture expansion and a growth arrest (GA) medium to model the proliferative quiescence of pericytes *in vivo*. Three media compositions **(Supplementary Table 1)** were tested at both hyperoxia **(Fig. 1B)** and physioxia **(Fig. 1C)** for the ability to induce pericyte proliferation or to maintain proliferative quiescence over 72 h. Medium 1 (M1) was a commercially available pericyte growth medium, medium 2 (M2) was an endothelial cell growth medium without additional growth factors, and medium 3 (M3) was the same endothelial cell growth medium supplemented with endothelial cell growth factors. The GA version of each medium composition had reduced serum and no additional growth factors. GM composition M3 supported pericyte culture expansion better than M1 and M2 under physioxia conditions. Additionally, the GA version of composition M3 effectively limited pericyte proliferation over the seeding density (indicated by the dashed line) in both oxygen tensions, maintaining pericytes in proliferative quiescence. The ability to effectively reduce proliferation accurately models pericyte quiescence associated with blood vessel investment under homeostatic conditions.

**Figure 1.**
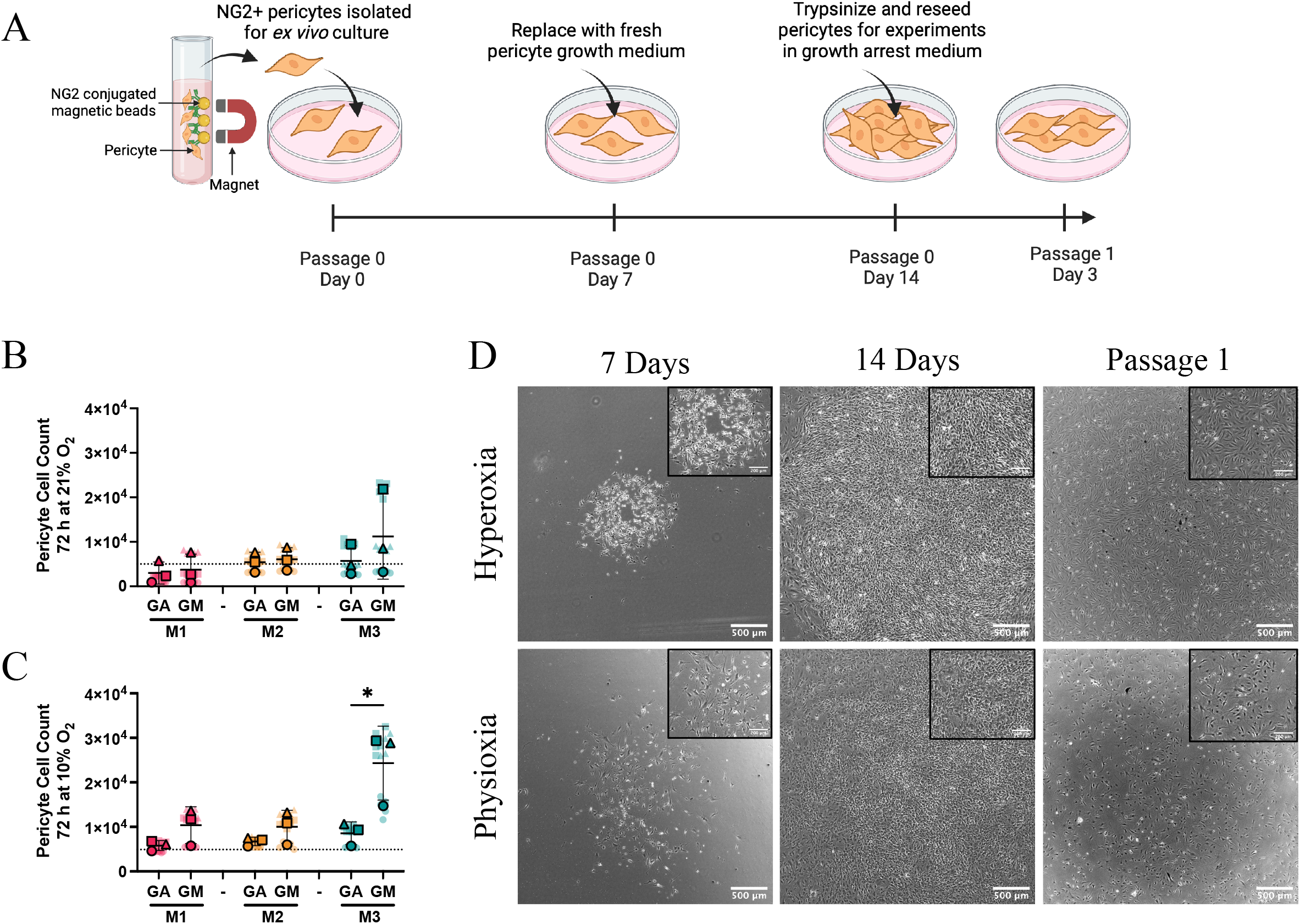
Physiological oxygen levels support primary lung pericyte expansion *ex vivo*. **(A)** Schematic demonstrating the timeline from pericyte isolation to use in experiments. **(B-C)** Pericyte proliferation over 72 h in response to both the growth arrest (GA) and growth medium (GM) compositions of the media to be optimised (M1-3). Proliferation at hyperoxia is displayed in **(B)** and lung physioxia in **(C)**. Dotted line represents seeding density. Statistical significance was assessed by unpaired t tests **(B-C)** (*p<0.05). Data are displayed as Superplots (Large symbols: means of experimental replicates; Small symbols: technical replicates for each experimental replicate; Lines: mean ±SD of replicate means). **(D)** Representative phase contrast images of primary lung pericytes expanded in medium composition M3 and maintained in either hyperoxia (21% O_2_) or lung physioxia (10% O_2_). Key time points during the expansion of primary pericytes are displayed. Schematic made in Biorender.com.

Phase contrast images were taken during the course of pericyte expansion to assess morphology in each medium. At 7 d post isolation, cells in physioxia and hyperoxia had visible pericyte colonies. However, only M3 achieved confluence by 14 d **(Fig. 1D)**, with M1 and M2 remaining as small colonies **(Supplemental Fig. 1)**. Pericytes maintained in M1 or M2 exhibited morphological signs of senescence after passage 1 in physioxia **(Supplemental Fig. 1)**. In contrast, pericytes from M3 maintained in physioxia retained typical pericyte morphology with multiple long cellular projections after passage 1. However, pericytes maintained in hyperoxia appeared fibroblastic, indicated by the spindle-shaped cell morphology and parallel alignment^27^ **(Fig. 1D)**. These data suggested that the oxygen tension under which pericytes are grown can influence phenotypic status, indicated by differences in cell morphology.

### Primary lung pericytes cultured in physioxia demonstrate stable transcript expression of characteristic genes

To assess the culture conditions that best-retained pericyte identity *ex vivo*, we analyzed changes in expression of *Cspg4* **(Fig. 2A)** and *Pdgfrb* **(Fig. 2B)**, pericyte identification markers, under physiologically relevant GA conditions with 10% O_2_ compared to 21% O_2_. Interestingly, expression of both genes was retained in all experimental conditions. Transcriptional expression of the characteristic perivascular marker genes, *Acta2* **(Fig. 2C)** and *Myh11* **(Fig. 2D)** were also retained *ex vivo* in all conditions. Variance in gene expression was calculated between independent pericyte batches and across culture conditions, demonstrating that composition M3 and physioxia promoted stability of transcriptional expression **(Fig. 2F)**.

**Figure 2.**
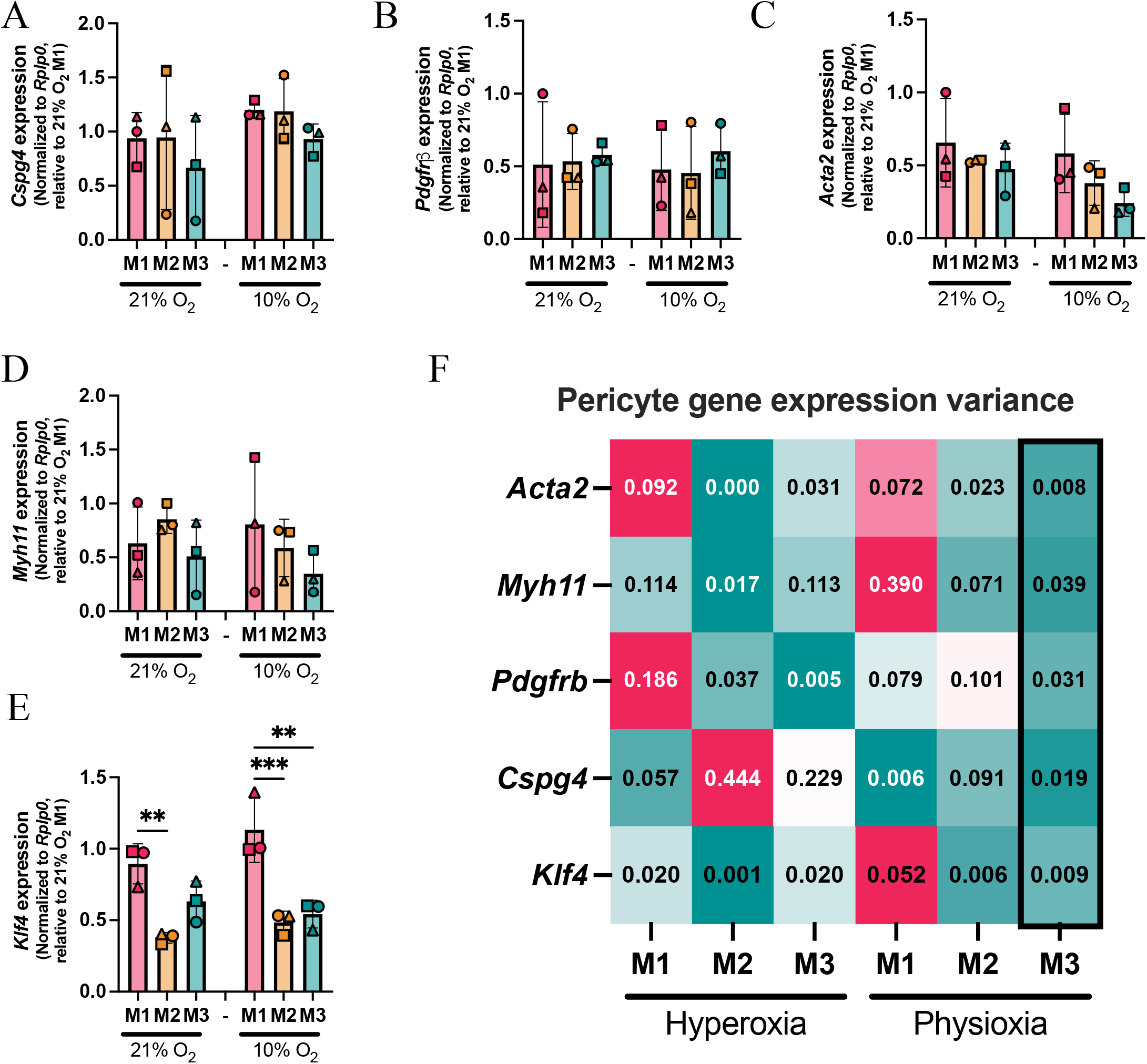
Physioxia reduces gene expression variation between pericyte batches. **(A)** RT-qPCR was performed on pericytes expanded in three different medium compositions under physioxia and hyperoxia to assess transcriptional expression of the characteristic pericyte genes *Cspg4* **(A)**, *Pdgfrb* **(B)**, *Acta2* **(C)**, *Myh11* **(D)** and *Klf4* **(E)**. Data are displayed as average fold change per experimental replicate ±SD (N=3). Statistical significance was assessed by one-way ANOVA with Tukey’s multiple comparisons test (*p<0.05, **p<0.01). Variance of pericyte gene expression data is plotted in the heat map **(F)**.

It is reported in pathologies, including cancer, that pericytes undergo phenotypic switching, indicated by increased expression of the pluripotency gene, *Klf4*^8^. Pericyte *Klf4* expression was observed to be significantly increased when maintained in medium composition M1 under both oxygen tensions **(Fig. 2E & 2F)** suggesting that it had an ‘activating’ effect on pericytes. Here we have identified the culturing conditions that retain characteristic pericyte identification and functional genes, while minimising the expression of the activating factor *Klf4*, to model physiologically relevant pericyte biology. The data suggest that medium composition M3 will aid reproducibility of experiments by limiting the gene expression variation between independent pericyte batches.

### Physioxic growth conditions aid culture of pericytes from multiple organ sources

Pericytes are found throughout the body and are highly heterogenous in function and morphology based on their location along capillary beds as well as between different organs. To test whether our protocol would be applicable to pericytes derived from multiple organ sites beyond the lung, NG2-expressing cells were isolated from brain, bone, and liver. *In vitro* cultures resembled typical pericyte morphology with a prominent cell body and multiple long extensions regardless of anatomic origin **(Fig. 3, Supplemental Fig. 2)**. Pericyte morphology was compared between hyperoxia (21% O_2_) and the appropriate physioxia for each organ-derived cell line (5% O_2_ for brain, bone, and liver; 10% O_2_ for lung).

**Figure 3.**
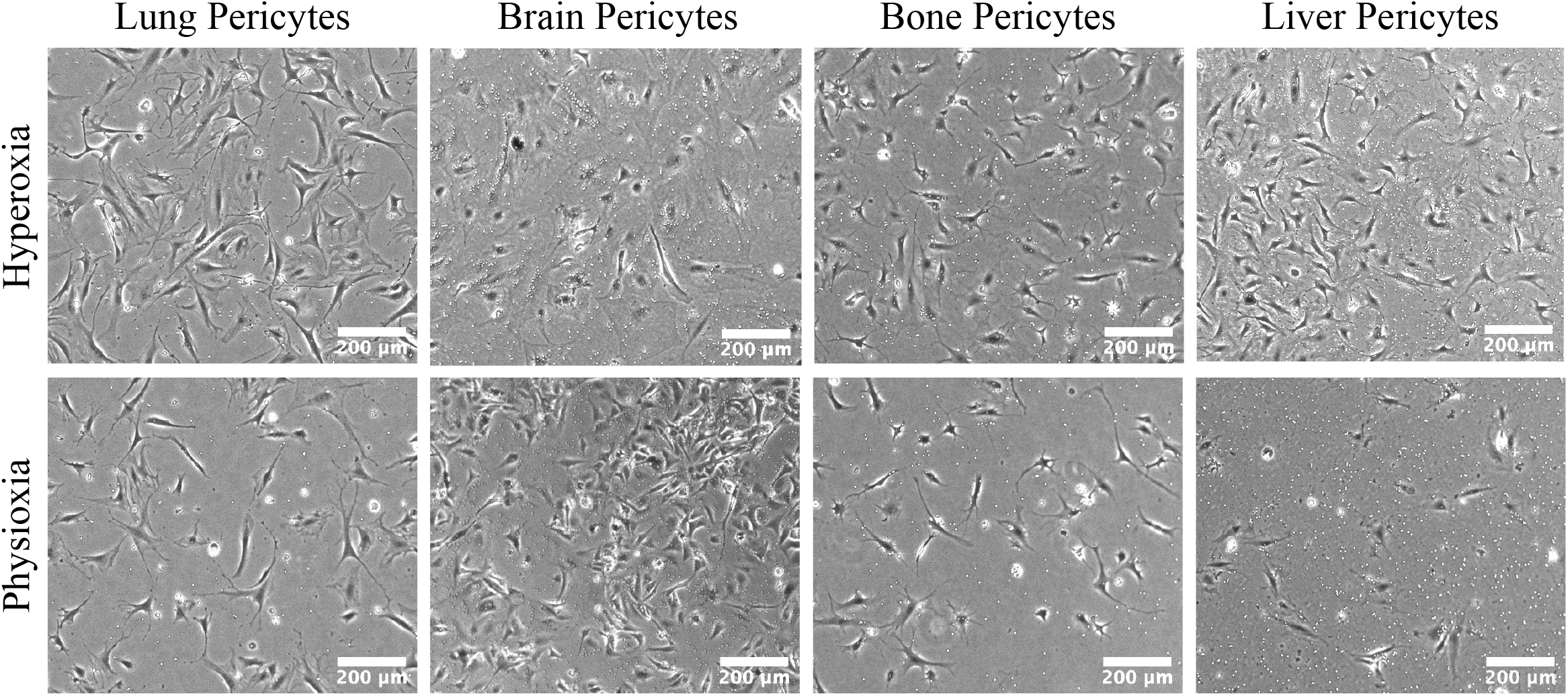
Primary pericytes expanded in physioxia retain characteristic morphology. Representative phase contrast images of primary lung, brain, bone, and liver pericytes expanded in M3 and maintained in either hyperoxia (21% O_2_) or physioxia (10% O_2_ for lung; 5% O_2_ for brain, bone and liver). Scale bar = 200 μm.

Protein expression of characteristic pericyte markers was validated across all organs using immunofluorescence for 4 key proteins, NG2, PDGFRb MYH11, and ACTA2 **(Fig. 4)**. A heterogeneous staining pattern was observed within the pericyte cultures, suggesting this culture method can capture the heterogeneity of pericyte phenotypes observed *in vivo* along the microvascular tree. Capturing multiple pericyte subtypes provides a more accurate representation of the *in vivo* pericyte population, which will improve the translational relevance of this model system.

**Figure 4.**
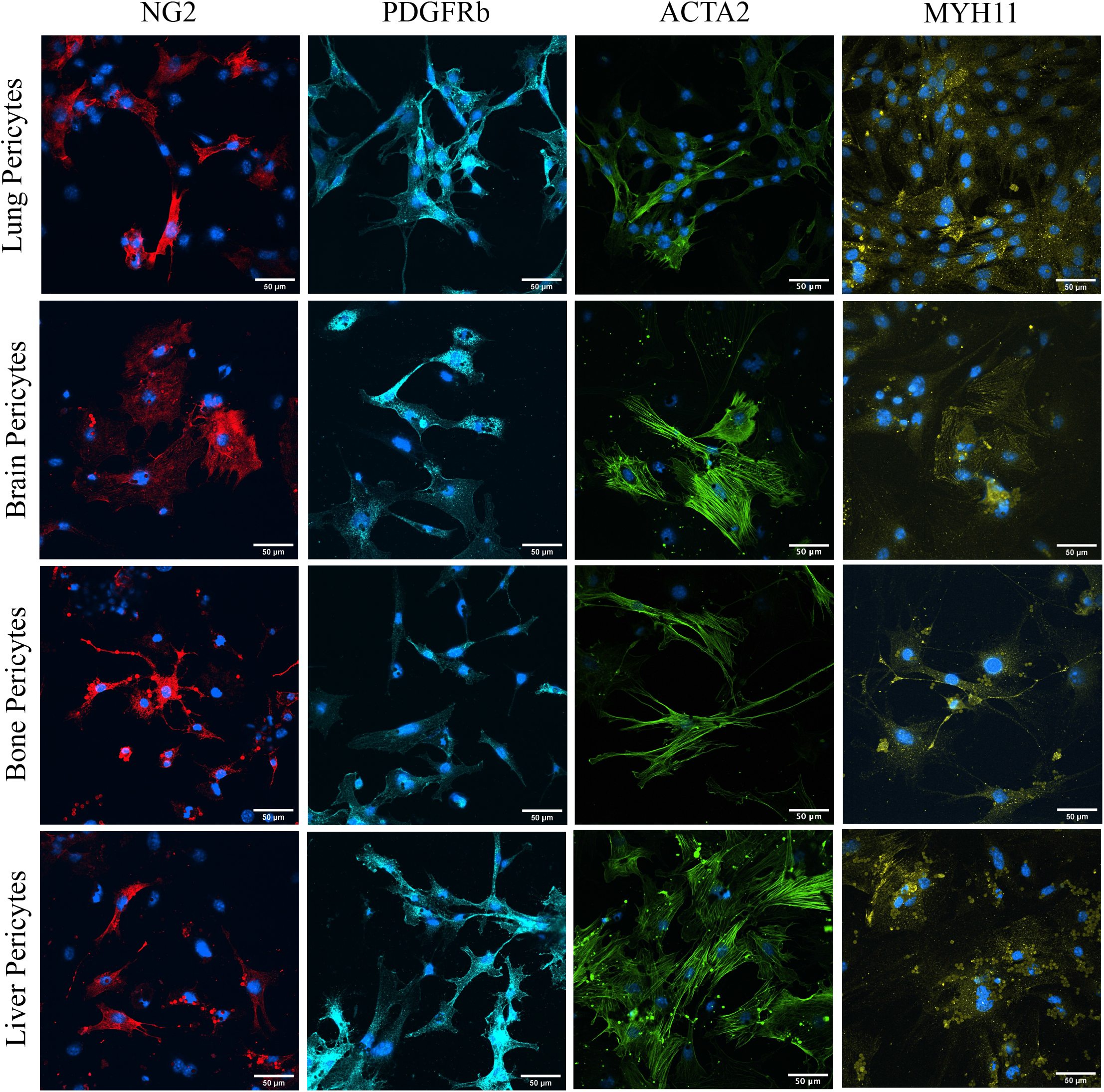
Immunofluorescence characterization of primary pericyte cultures from mouse. Pericytes obtained from mouse lung, brain, bone, and liver were seeded at passage 1 for immunofluorescence study. A panel of pericyte identification markers including NG2, MYH11, PDGFRb, and ACTA2 were used to characterize pericyte marker expression in different organs. One representative image is displayed from a total of 3 independent pericyte batches. Scale Bar = 50 μm.

### *Ex vivo* pericyte cultures demonstrate a functionally relevant association with endothelial junctions

A key physiological role of pericytes is to support blood vessel formation and structural integrity. To determine if pericytes retained their physiological function in vascular support *ex vivo*, tube formation assays were performed using human umbilical vein endothelial cells (hUVECs) in co-culture with zsGreen expressing pericytes. Pericyte density on vessels is known to correlate with the barrier function unique to each organ^28^. Pericyte density is highest in the central nervous system (CNS) with up to 100% vessel coverage^29,30^ compared to the lungs and liver, having an approximately 1:10 ratio of pericytes to endothelial cells (ECs)^28^. To account for these differences in pericyte investment, tube formations were performed with pericytes from lung, bone, and liver co-cultures used at a 1:10 ratio while brain pericytes were co-cultured at a 1:5 ratio of pericyte to ECs. hUVECs demonstrated alignment and adhesion in Matrigel to form tube-like structures. Pericytes isolated from the lung, brain, bone, and liver when added to co-culture associated with the endothelial tubes. Representative images are shown for the 4 h time point in each culture condition (**Fig. 5A**). The tube formation images were analyzed to determine the localization of pericytes to endothelial tubes and were assigned junction, tube, or unassociated **(Supplemental Fig. 3)**. Pericytes from all organs were significantly associated with endothelial junctions (**Fig. 5B-E**). These data support known pericyte physiological function and location on branch points in the vasculature^31^, indicating that these pericyte cultures retain physiologically relevant functions.

**Figure 5.**
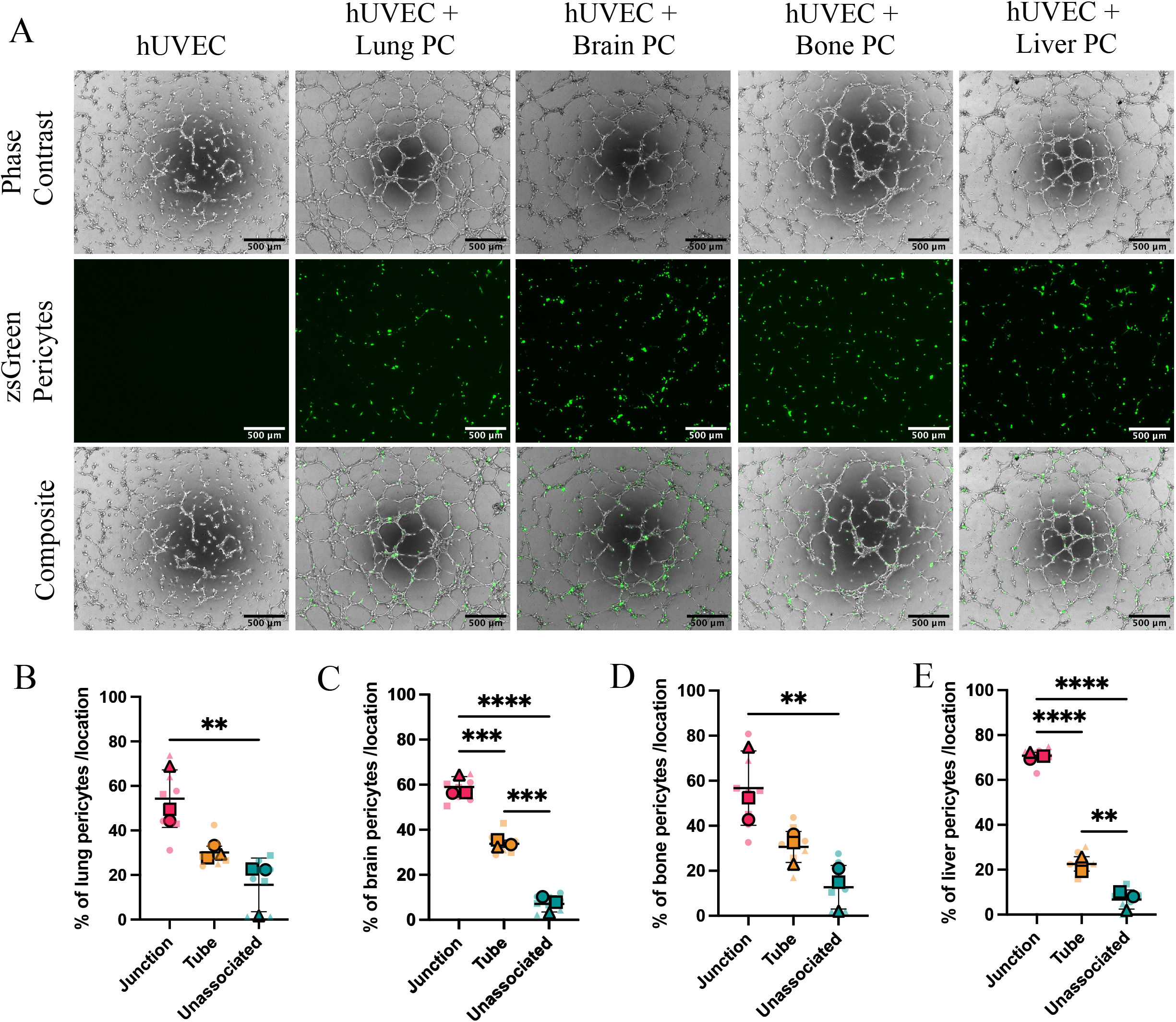
*Ex vivo* pericyte retain physiological function at endothelial junctions. hUVECs (unlabeled) were co-cultured with pericytes (zsGreen) at a ratio of 1:10 (lung, bone, and liver) or 1:5 (brain) PC:EC in Matrigel over 4 h. Representative images from 4 h co-culture are displayed **(A)**. Scale Bar = 500 µm. Quantification of pericyte association with endothelial tubes is quantified as the number of zsGreen positive cells found at junctions, tubes, or unassociated **(B-E)**. Statistical significance was assessed by one-way ANOVA with Tukey’s multiple comparison test (**p<0.01, ***p<0.001, ****p<0.0001). Data are displayed as Superplots (Large symbols: means of experimental replicates; Small symbols: technical replicates for each experimental replicate; Lines: mean ±SD of replicate means).

### Primary lung pericytes cultured in physioxia accurately model tumor-derived factor induced activation

Our previous work has demonstrated that metastatic tumor-derived factors promote pericyte activation, characterized by pericyte proliferation, down-regulation of perivascular marker genes, and an increase in *Klf4* expression^8^. To examine whether *ex vivo* pericyte cultures recapitulate our previous *in vivo* findings, lung pericytes were treated with tumor cell conditioned medium (TCM) collected from the metastatic triple-negative breast cancer cell line, 4T1. Pericyte activation conditions were pre-determined by assessing proliferation in response to different concentrations of TCM and monitoring *Klf4* expression over time **(Supplemental Fig. 4)**. Pericytes were treated with 50% TCM for 72 h, and proliferation was assessed compared to the GA control. 4T1 TCM induced significant pericyte proliferation **(Fig. 6A)**. The expression of the pluripotency factor, *Klf4*, was also significantly increased in pericytes 30 min after exposure to 4T1 TCM compared to the GA control **(Fig. 6B)**. Additionally, perivascular cell marker genes, *Myh11* and *Acta2* were significantly downregulated in pericytes treated with 4T1 TCM after 72 h compared to GA **(Fig. 6C-D)**. These parameters together indicate that 4T1 TCM induced pericyte phenotypic switching like that observed *in vivo*, suggesting that these pericyte cultures can be used to study the role of pericytes in metastasis.

**Figure 6.**
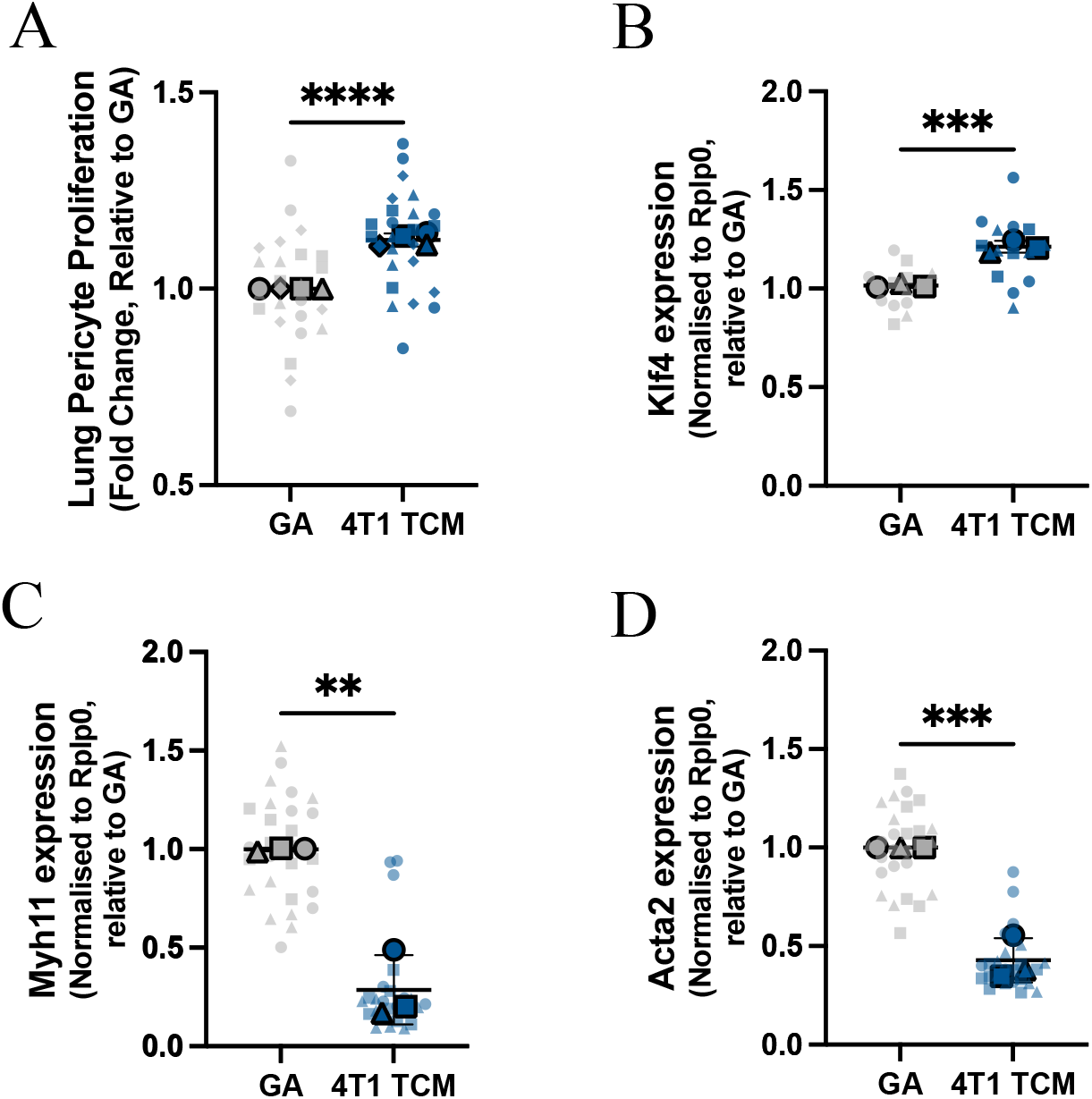
*Ex vivo* pericyte cultures accurately recapitulate *in vivo* phenotypic switching. **(A)** Pericyte phenotypic switching *ex vivo* was assessed by treating lung pericytes with TCM from the metastatic breast cancer cell line 4T1. Lung pericyte proliferation in response to TCM was assessed at 72 h **(A)** and *Klf4* expression was assessed by RT-qPCR at 30 min following TCM treatment **(B)**. Expression of characteristic perivascular marker genes, *Myh11* **(C)** and *Acta2* **(D)** were assessed after 72 h of TCM treatment compared to the GA control. Statistical significance was assessed by unpaired t tests (**p<0.01, ***p<0.001, ****p<0.0001). Data are displayed as Superplots (Large symbols: means of experimental replicates; Small symbols: technical replicates for each experimental replicate; Lines: mean ±SD of replicate means).

## Discussion

Historically, it has been difficult to model pericyte biology *in vitro* due to low yield and lack of standardized methodologies^3^. Limited pericyte cell lines are available and major problems are associated with the current tools. Under homeostatic conditions pericytes are quiescent and not actively proliferating unless required in angiogenesis or response to injury^32^ whereas immortalised cell lines are induced to permanently proliferate^33^, a phenotype associated with perivascular cell activation^8,34^. Pericytes have been generated successfully from induced pluripotent stem cells (iPSC)^35–37^ however iPSC can be difficult to produce and require highly specific culturing conditions^38,39^. Primary cell lines are a useful tool to recapitulate *in vivo* variation and physiology but are limited by expansion capabilities and the retention of cellular function and phenotype *ex vivo*^3,40^. These challenges have resulted in pericytes remaining an understudied population despite their demonstratably vital functions in both physiology and disease.

In this study we present an NG2-specific antibody conjugated magnetic bead pull down protocol for the isolation of pericytes from multiple organ sources. We focused on NG2-expressing pericytes because we have previous found that NG2+ pericytes promote cancer metastasis^8^. Additionally, NG2 appears to be expressed continuously on arteriole and mid-capillary pericytes and therefore should capture a range of known pericyte subtypes^41^; however, in addition or alternatively to NG2, pericytes can express proteins such as PDGFRb^3,42^. Future studies may highlight whether a similar approach could be used for alternative pericyte subpopulations including PDGFRb-expressing pericytes which may not all express NG2.

Optimization of pericyte culturing conditions highlighted the importance of maintaining primary cultures in physioxia, as oxygen tension is becoming increasingly appreciated as a mediator of cellular phenotypic switching^43,44^. For example, physioxia was found to be critical in maintaining hepatocyte morphology and function *ex vivo*, with cultures maintained at hyperoxia rapidly undergoing epithelial-mesenchymal transition (EMT) and losing hepatocyte functions including albumin production, low-density lipoprotein (LDL) uptake, and expression of drug metabolizing enzymes^22^. We have previously found that culturing microvascular endothelial cells (ECs) at hyperoxia impaired the cellular response to hypoxia, when compared to those cells that had been maintained in oxygen tensions that matched those found in their tissue of origin^21^. Consistent with these reports, pericyte maintenance in physioxia better recapitulated pericyte morphology, characteristic marker expression and association with endothelial tubes.

Pericytes exist as a continuum along microvascular arterioles to venules and have unique functional roles depending on location^1^, with ensheathing pericytes acting to rapidly control blood flow, and thin-strand pericytes functioning as metabolic sensors^24,45^. Additionally, each organ has a specialized microvascular network to carry out distinct functions. Continuous capillaries, which are found in most organs including the brain and retina, are barrier forming, whereas sinusoidal capillaries, found in the liver and bone marrow, are discontinuous and allow free movement of materials^46^. Recent studies have noted that pericytes analyzed from multiple organs retain expression of organ-specific solute carriers critical for their organ-specific function^47,48^. Importantly, we found that pericytes isolated from lung, brain, bone and liver displayed heterogeneity in morphology and expression of functional proteins suggesting that this method preserves pericyte function and organ specificity.

The overall goal of our study was to create a pericyte culture model that could be used to better understand the role of pericytes in metastasis. Critical to the success of creating an *in vitro* primary pericytes model system was the ability to recapitulate lung pericyte activation during metastatic progression, as was observed *in vivo*^8^. We found that primary lung pericytes expressed the stem/plasticity factor *Klf4* and increased proliferation in response to TCM from the metastatic breast cancer cell line, 4T1. Additionally, pericytes treated with 4T1 TCM also demonstrated reduced expression of characteristic perivascular marker genes *Myh11* and *Acta2*, suggesting that pathological phenotypic switching can be observed in thi*s ex vivo* system. These pericyte responses are hallmarks of their plasticity and activation during premetastatic niche formation^8^.

In this study, we have described a high throughput and reproducible pericyte cell culture system that is dependent on physiologically relevant oxygen tensions to accurately recapitulate pericyte biology. This tool will aid the study of pericytes in health and disease, including investigation into the organ specific roles of pericytes in metastasis.

## Materials and Methods

### Animals

Pericyte cell cultures were obtained from female C57BL/6 mice at 12 weeks of age. Animals were anesthetized by isoflurane followed by cervical dislocation. All animals were maintained under pathogen free conditions within the NIH animal facility. All animal experiments were conducted in accordance with the guidelines and regulations outlined in the Guide for the Care and Use of Laboratory Animals and were approved by the NCI-Bethesda Animal Care and Use Committee. All procedures were in accordance with those outlined in the ARRIVE guidelines.

### Primary pericyte isolation and culture

A protocol for the isolation of murine pericytes was developed by adapting a previously published protocol for the isolation of microvascular ECs^21,25^. The pericyte isolation protocol has been described in detail in supplemental methods and materials **(Supplemental Fig. 5)**. Pericyte yield and behavior may be altered by many factors including age, gender or pathology, therefore, to standardize the protocol only 12-week-old female C57BL/6 mice were used for pericyte cultures. Pericytes were cultured on plates coated with 0.06 mg/mL collagen (C7661, Millipore Sigma) and maintained in a 37°C humidity-controlled incubator at physioxia (10% O_2_ for lungs or 5% O_2_ for bone, brain, and liver, with 5% CO_2_) and used for experiments within 10-14 d. Pericytes were maintained in medium containing a 1:1 mix of low glucose Dulbecco’s Modified Eagle’s medium (DMEM) (D6046, Millipore Sigma) and Ham’s F12 nutrient mix (11765047, Thermo Fisher Scientific), supplemented with 1% MEM non-essential amino acids (11140050, Thermo Fisher Scientific), 2mM sodium pyruvate (11360070, Thermo fisher Scientific), 20mM Hepes (15630080, Thermo Fisher Scientific), 1X endothelial cell growth factors (390599, Bio-techne), 1% penicillin/streptomycin (15070063, Thermo Fisher Scientific) and 20% fetal bovine serum (900-208-500, GeminiBio). Before experiments pericytes were placed in GA medium containing reduced FBS (2%) and no endothelial cell growth factors for 24 h.

### Cell lines and culturing conditions

Human umbilical vein endothelial cells (hUVEC) were provided by Clare Waterman (NHLBI, Bethesda, USA) and cultured on plastic in a 37°C humidity-controlled incubator with 5% CO_2_. medium containing a 1:1 mix of low glucose DMEM and Ham’s F12 nutrient mix, supplemented with 1% MEM non-essential amino acids, 2mM sodium pyruvate, 20mM Hepes, 2% endothelial cell growth factors, 10 mg/mL heparin (H3149, Millipore Sigma), 1% penicillin/streptomycin and 10% fetal bovine serum.

The metastatic murine 4T1 tumor cell line was provided by Kent Hunter (National Cancer Institute, Bethesda, USA). The 4T1 cell line was maintained on 2D plastic culture flasks in a 37°C humidity-controlled incubator with 5% CO_2_ in RPMI (11-875-119, Thermo Fisher Scientific) with 10% FBS, 1% penicillin/streptomycin.

### Tumor cell conditioned media collection

Tumor cells were plated at 1×10^5^ cells/mL in a T75 flask with pericyte GA medium and allowed to proliferate for 72 h. The tumor cell conditioned medium (TCM) was then collected and filtered through a 0.22 μm syringe filter (380111, Nest Scientific), aliquoted and stored at -80°C for downstream experiments. The TCM stock was diluted 1:1 in pericyte GA medium before use in experiments.

### Proliferation assay

Pericytes were seeded on collagen coated 96-well plates at 5×10^3^ cells per well for lung pericytes and 1×10^4^ cells per well for brain, bone, and liver pericytes in GA medium and returned to the appropriate incubator to adhere overnight. Pericytes were treated with control GA medium or TCM from 4T1 tumor cells that was diluted 1:1 in GA medium. The treated cells were returned to the appropriate incubator, 72 h later the cell nuclei were stained with 2 µM Hoechst (62249, Thermo Fisher Scientific) in PBS and the cells were counted using the Celigo image cytometer.

### Quantitative real-time PCR

For optimisation of pericyte growth conditions RNA was collected from pericytes seeded at 1×10^5^ in a 12-well plate using the RNeasy Mini Kit (74104, Qiagen) after 24 h in the appropriate GA medium. For pericyte phenotypic switching experiments, pericytes were seeded in GA medium and allowed to adhere overnight, the following day the cells were treated with 4T1 TCM or GA medium for 72 h, and then isolated as above. RNA was quantified using a Nanodrop and 200 ng was used for reverse transcriptase cDNA synthesis using the iScript Reverse Transcriptase Supermix (1708841, BioRad). RT-qPCR was performed using cDNA diluted 1:30 in RNA free water, EvaGreen master mix (1725212, BioRad) and 10 µM primers.

Alternatively, for 30 min collections lung pericytes were seeded on 96-well plates with 5×10^3^ cells per well. The pericytes were seeded in GA medium and allowed to adhere overnight in the appropriate incubator. The following day the pericytes were treated with 4T1 TCM or GA as a control. After 30 min the treatments were removed, the cells were washed once with ice cold PBS, and collected immediately using the cell-2-cDNA kit (AM1723, Invitrogen). Multiplex qPCR was performed with cDNA samples diluted 1:4 and IDT prime time master mix (2x) (1055771, IDT). Primers for house-keeping gene *Rplp0* were used as a concentration of 250 nM each, while primers for *Klf4* were used at 500 nM, probes used at 500 nM.

All primers were designed using IDT and sequences can be found in **Supplementary Table 2**. Analysis was performed using the ΔΔCt method, using *Rplp0* as the reference gene, and data is presented as average fold change ± standard deviation (SD).

### Immunofluorescence

Millicell EZ 8-Well chamber slides (PEZGS0816, Millipore Sigma) were prepared with 0.06 mg/mL collagen, for a minimum of 2 h at 37°C. The collagen was removed from the wells and rinsed once with sterile PBS prior to seeding the cells. Pericytes were seeded at 3×10^4^ cells per chamber in 150 µL GA medium and allowed to adhere overnight. The next day the medium was removed from the chamber slides and the cells were fixed with ice cold 70% methanol for 7 min. The cells were permeabilised with 0.5% Triton X-100, blocked with 10% donkey serum (S30-M, Millipore Sigma)/3% Fish Gelatin (G7765, Sigma Aldrich) and incubated with primary antibodies at 4°C overnight. The following primary antibodies were used; rabbit anti-MYH11 (ab224804, Abcam), rabbit anti-PDGFRb (ab32570, Abcam), recombinant Alexa Fluor 647 anti-NG2 (ab283639, Abcam), and mouse anti-ACTA2 FITC conjugated (F3777, Millipore Sigma). The following day the cells were washed with 0.1% Tween20 (655204, Millipore Sigma) before incubation with donkey secondary antibodies (Jackson ImmunoResearch) for 1 h at room temperature, protected from light. The cells were washed again in 0.1% Tween20 solution followed by incubation with DAPI (D1306, Thermo Fisher Scientific) and finally mounted with ProLong™ Gold Antifade mountant (P36930, Thermo Fisher Scientific). Images were obtained on the Nikon SoRa spinning disk microscope.

### Tube formation assays

10 µL of ice cold Matrigel (3432-001-01, R&D) was loaded onto 15 µ-angiogenesis slide (81506, Ibidi) and Matrigel was allowed to polymerise for 30 min at 37°C. hUVECs were trypsinised, centrifuged and washed in endothelial medium without FBS or growth factors. hUVECs were seeded at 1×10^4^ cells per well in 50 µl of serum free medium onto the Matrigel. For pericyte-EC tube formation assays, pericytes stably expressing the zsGreen fluorescent protein were added to the hUVECs at a ratio of 1:5 for brain pericytes or 1:10 for lung, bone and liver pericytes (Pericyte: EC). Tube formation was imaged every 30 min for 8 h using the BioTek Lionheart FX automated microscope (LFXW-SN, Agilent).

### Statistical Analysis

Data are presented as mean ± SD. Where relevant, statistics were performed on the mean values of the independent experiments. Statistical tests were performed using GraphPad Prism 5 software and statistical significance was accepted at the 95% confidence level. Image analysis was performed using ImageJ ^49^.

## Data Availability Statement

All data generated or analyzed during this study are included in this article and Supplementary Information files.

## Author Contributions

TM, ECM, MJG, and LEZ conducted the experiments and analysed the data. TM prepared the manuscript draft. MM and CMB designed and supervised the project. All authors reviewed and approved the final version of the manuscript.

## References

1. Holm, A., Heumann, T. & Augustin, H. G. Microvascular Mural Cell Organotypic Heterogeneity and Functional Plasticity. Trends Cell Biol. 28, 302–316 (2018).

2. Armulik, A., Genové, G. & Betsholtz, C. Pericytes: Developmental, Physiological, and Pathological Perspectives, Problems, and Promises. Dev. Cell 21, 193–215 (2011).

3. Alvino, V. V., Mohammed, K. A. K., Gu, Y. & Madeddu, P. Approaches for the isolation and long-term expansion of pericytes from human and animal tissues. Frontiers Cardiovasc Medicine 9, 1095141 (2023).

4. Splunder, H. van, Villacampa, P., Martínez-Romero, A. & Graupera, M. Pericytes in the disease spotlight. Trends Cell Biol. (2023) doi:10.1016/j.tcb.2023.06.001.

5. Hosaka, K. et al. Pericyte–fibroblast transition promotes tumor growth and metastasis. Proc. Natl. Acad. Sci. 113, E5618–E5627 (2016).

6. Yamaguchi, M. et al. Pericyte-myofibroblast transition in the human lung. Biochem. Biophys. Res. Commun. 528, 269–275 (2020).

7. Nobre, A. R. et al. Bone marrow NG2+/Nestin+ mesenchymal stem cells drive DTC dormancy via TGF-β2. Nat. Cancer 2, 327–339 (2021).

8. Murgai, M. et al. KLF4-dependent perivascular cell plasticity mediates pre-metastatic niche formation and metastasis. Nat Med 23, 1176–1190 (2017).

9. Prazeres, P. H. D. M. et al. Pericytes are heterogeneous in their origin within the same tissue. Dev Biol 427, 6–11 (2017).

10. Yamazaki, T. & Mukouyama, Y. Tissue Specific Origin, Development, and Pathological Perspectives of Pericytes. Front. Cardiovasc. Med. 5, 78 (2018).

11. Hartmann, D. A., Coelho-Santos, V. & Shih, A. Y. Pericyte Control of Blood Flow Across Microvascular Zones in the Central Nervous System. Annu. Rev. Physiol. 84, 1–24 (2021).

12. Crisan, M. et al. A Perivascular Origin for Mesenchymal Stem Cells in Multiple Human Organs. Cell Stem Cell 3, 301–313 (2008).

13. Birbrair, A. et al. How Plastic Are Pericytes? Stem Cells Dev. 26, 1013–1019 (2017).

14. Birbrair, A. et al. Role of Pericytes in Skeletal Muscle Regeneration and Fat Accumulation. Stem Cells Dev. 22, 2298–2314 (2013).

15. Chen, C. et al. Physioxia: a more effective approach for culturing human adipose-derived stem cells for cell transplantation. Stem Cell Res. Ther. 9, 148 (2018).

16. D’Ippolito, G., Diabira, S., Howard, G. A., Roos, B. A. & Schiller, P. C. Low oxygen tension inhibits osteogenic differentiation and enhances stemness of human MIAMI cells. Bone 39, 513–522 (2006).

17. Holzwarth, C. et al. Low physiologic oxygen tensions reduce proliferation and differentiation of human multipotent mesenchymal stromal cells. BMC Cell Biol. 11, 11 (2010).

18. Kim, D. S. et al. Effect of low oxygen tension on the biological characteristics of human bone marrow mesenchymal stem cells. Cell Stress Chaperones 21, 1089–1099 (2016).

19. Adebayo, A. K. & Nakshatri, H. Modeling Preclinical Cancer Studies under Physioxia to Enhance Clinical Translation. Cancer Res. 82, 4313–4321 (2022).

20. McKeown, S. R. Defining normoxia, physoxia and hypoxia in tumours—implications for treatment response. Br. J. Radiol. 87, 20130676 (2014).

21. Reiterer, M., Eakin, A., Johnson, R. S. & Branco, C. M. Hyperoxia Reprogrammes Microvascular Endothelial Cell Response to Hypoxia in an Organ-Specific Manner. Cells 11, 2469 (2022).

22. Guo, R., Xu, X., Lu, Y. & Xie, X. Physiological oxygen tension reduces hepatocyte dedifferentiation in in vitro culture. Sci. Rep. 7, 5923 (2017).

23. Hirunpattarasilp, C. et al. Hyperoxia evokes pericyte-mediated capillary constriction. J. Cereb. Blood Flow Metab. 42, 2032–2047 (2022).

24. Hariharan, A., Robertson, C. D., Garcia, D. C. G. & Longden, T. A. Brain capillary pericytes are metabolic sentinels that control blood flow through a KATP channel-dependent energy switch. Cell Rep. 41, 111872 (2022).

25. Branco-Price, C. et al. Endothelial Cell HIF-1α and HIF-2α Differentially Regulate Metastatic Success. Cancer Cell 21, 52–65 (2012).

26. Reiterer, M. et al. Acute and chronic hypoxia differentially predispose lungs for metastases. Sci. Rep. 9, 10246 (2019).

27. Tomasek, J. J. & Hay, E. D. Analysis of the role of microfilaments and microtubules in acquisition of bipolarity and elongation of fibroblasts in hydrated collagen gels. J. cell Biol. 99, 536–549 (1984).

28. Zhang, Z.-S. et al. Research advances in pericyte function and their roles in diseases. Chin. J. Traumatol. 23, 89–95 (2020).

29. Mathiisen, T. M., Lehre, K. P., Danbolt, N. C. & Ottersen, O. P. The perivascular astroglial sheath provides a complete covering of the brain microvessels: An electron microscopic 3D reconstruction. Glia 58, 1094–1103 (2010).

30. Armulik, A. et al. Pericytes regulate the blood–brain barrier. Nature 468, 557–561 (2010).

31. Bergers, G. & Song, S. The role of pericytes in blood-vessel formation and maintenance. Neuro-oncology 7, 452–464 (2005).

32. Nwadozi, E., Rudnicki, M. & Haas, T. L. Metabolic Coordination of Pericyte Phenotypes: Therapeutic Implications. Front. Cell Dev. Biol. 8, 77 (2020).

33. Li, P. et al. Generation of a new immortalized human lung pericyte cell line: a promising tool for human lung pericyte studies. Lab Invest 101, 625–635 (2021).

34. Dzau, V. J., Braun-Dullaeus, R. C. & Sedding, D. G. Vascular proliferation and atherosclerosis: New perspectives and therapeutic strategies. Nat. Med. 8, 1249–1256 (2002).

35. Faal, T. et al. Induction of Mesoderm and Neural Crest-Derived Pericytes from Human Pluripotent Stem Cells to Study Blood-Brain Barrier Interactions. Stem Cell Rep. 12, 451–460 (2019).

36. Szepes, M. et al. Dual Function of iPSC-Derived Pericyte-Like Cells in Vascularization and Fibrosis-Related Cardiac Tissue Remodeling In Vitro. Int. J. Mol. Sci. 21, 8947 (2020).

37. Stebbins, M. J. et al. Human pluripotent stem cell-derived brain pericyte-like cells induce blood-brain barrier properties. bioRxiv 387100 (2018) doi:10.1101/387100.

38. Ghaedi, M. & Niklason, L. E. Organoids, Stem Cells, Structure, and Function. Methods Mol. Biol. 1576, 55–92 (2016).

39. Santostefano, K. E. et al. A practical guide to induced pluripotent stem cell research using patient samples. Lab. Investig. 95, 4–13 (2015).

40. Lee, L. L., Khakoo, A. Y. & Chintalgattu, V. Isolation and Purification of Murine Cardiac Pericytes. J Vis Exp (2019) doi:10.3791/59571.

41. Murfee, W. L., Skalak, T. C. & Peirce, S. M. Differential Arterial/Venous Expression of NG2 Proteoglycan in Perivascular Cells Along Microvessels: Identifying a Venule-Specific Phenotype. Microcirculation 12, 151–160 (2005).

42. Alarcon-Martinez, L., Yemisci, M. & Dalkara, T. Pericyte morphology and function. Histol. Histopathol. 36, 633–643 (2021).

43. Mattia, M. D. et al. Insight into Hypoxia Stemness Control. Cells 10, 2161 (2021).

44. Mas-Bargues, C. et al. Relevance of Oxygen Concentration in Stem Cell Culture for Regenerative Medicine. Int. J. Mol. Sci. 20, 1195 (2019).

45. Hartmann, D. A. et al. Brain capillary pericytes exert a substantial but slow influence on blood flow. Nat. Neurosci. 24, 633–645 (2021).

46. Augustin, H. G. & Koh, G. Y. Organotypic vasculature: From descriptive heterogeneity to functional pathophysiology. Science 357, (2017).

47. Baek, S.-H. et al. Single Cell Transcriptomic Analysis Reveals Organ Specific Pericyte Markers and Identities. Frontiers Cardiovasc Medicine 9, 876591 (2022).

48. Vanlandewijck, M. et al. A molecular atlas of cell types and zonation in the brain vasculature. Nature 554, 475–480 (2018).

49. Schneider, C. A., Rasband, W. S. & Eliceiri, K. W. NIH Image to ImageJ: 25 years of image analysis. Nat. Methods 9, 671–675 (2012).

