## Supplemental Figures for "Pericytes require physiological oxygen tension to maintain phenotypic fidelity"

### Supplementary Materials and Methods

#### Pericyte Isolation Protocol

**A. Preparation of materials and culture medium** (See **Supplementary Table 3** for a list of all materials required)

1. Autoclave micro dissecting scissors and forceps.
2. Warm Dulbecco's Modified Eagle's Medium – low glucose (DMEM-LG) to 37°C (required for brain digestion protocol).
3. Make complete growth medium and equilibrate in the relevant oxygen tension incubator for at least 24 h.
4. For brain digestions make a solution of 20% bovine serum albumin (BSA) in DMEM-LG. BSA will dissolve on a roller at 37°C. Once dissolved, filter the solution through a 0.2 µm filter. This solution may be aliquoted and stored at -20°C. 10 mL is needed per batch (typically 3-6 organs per batch).
5. Make phosphate buffer saline (PBS)/0.1% BSA and store at 4°C.
6. Collagen stock should be prepared in 0.1 M acetic acid at a concentration of 3 mg/mL. The collagen in acetic acid should be placed on a roller at room temperature for 2-3 h, followed by overnight rolling at 4°C. Once dissolved store the collagen at 4°C. Before use dilute the 3 mg/mL collagen 1:50 in sterile deionised (DI) water (final concentration 0.06 mg/mL). To coat the plates, pipette enough collagen per well to cover the surface of the well or flask and incubate for ≥ 2 h at 37°C.

**B. Preparation of Animal and Tissue Collection**

1. For each batch of pericytes, collect the organs from 3-6 animals. The data in this study were obtained from 12-week-old female C57BL/6 mice. However, this protocol should be adaptable to variables such as mouse age, sex or genetic background, as needed.
2. Anesthetise the mice with isoflurane and then perform cervical dislocation. Do not use $CO_2$ to euthanise the animals as this will alter pericyte behaviour.
3. Position the mouse in dorsal recumbency on a dissection board and use pins to secure the limbs extended in a diagonal position away from the body. Spray the mouse down with 70% ethanol.
4. Use forceps to grasp the skin over the sternum and make an incision with the scissors at the ventral midline. Extend the incision the length of the ventral midline cutting through the diaphragm, sternum, and ribs.
5. Make a small incision in the right atrium and cut the abdominal aorta below the liver.
6. Perfuse the mouse with 10 mL of ice-cold PBS containing 10 µg/mL heparin through the left ventricle, you should be able to visualise the organs becoming lighter in colour.

7. To extract the lungs, use the forceps to grab the trachea and cut to release the lungs, gently cut any connective tissue from behind as you go. Remove the heart from the lung block. Place the lungs in ice-cold DMEM-LG.
8. To extract the liver, use scissors to cut the liver away from the diaphragm and the abdominal organs. Once the liver is released, place it on a clean paper towel and identify the medial lobe that has a butterfly like appearance with the gall bladder in the middle. Gently clamp the end of the gall bladder closest to the tissue with your forceps and cut it away with scissors ensuring the gall bladder contents are not released over the tissue. Place the liver in ice-cold DMEM-LG.
9. To extract the femurs, extend the midline excision in the lower abdomen above the hip, down the leg to the ankle joint. Use the scissors to cut the anterior quadricep muscle exposing the femur. Position the scissors posterior to the femur to cut the hamstring at the knee joint. Place the scissor blades on either side of the femur and gently move them up to you reach the femoral head hip joint. With some pressure but without cutting, twist the scissors back and forth gently to dislocate the femur. Use forceps to hold the femur and cut away any remaining muscle. Use the scissors again at the knee joint to gently dislocate the femur from the tibia. Place the femur in ice-cold DMEM-LG.
10. To extract the brain, decapitate the mouse using scissors. Cut the skin on the dorsal midline and pull away the skin to expose the full skull. Insert one scissor blade into the spinal cord space and carefully cut along the sagittal suture to remove the calvaria (skull cap). Use forceps in a closed position to gently scoop under the brain, releasing any attachments and freeing the brain. Place the brain in ice-cold DMEM-LG.
11. Tissues may be stored on ice at 4°C for up to 12 h.

### C. Preparing the Dynabeads

1. Place a 1.5 mL microcentrifuge tube into a non-magnetic holder.
2. Swirl the bottle containing the Sheep Anti-Rat IgG Dynabeads immediately before use to ensure they are in suspension.
3. Pipette 10 µL of beads per organ collected and place this in the microcentrifuge tube in the non-magnetic holder.
4. Add 1 mL of PBS/0.1% BSA to the aliquot of beads and pipette up and down to ensure sufficient washing.
5. Move the microcentrifuge tube to the magnetic holder and allow the solution to turn clear (30-60 s).
6. Gently remove the PBS/0.1% BSA without disturbing the beads.
7. Repeat this washing step three times, then leave the beads in the fourth 1 mL of PBS/0.1% BSA in a non-magnetic holder.
8. Add 1.2 µL of the NG2 monoclonal antibody per organ to the washed bead solution. e.g., for five lungs, add 6 µL of antibody to 50 µL of beads in 1 mL of PBS/0.1% BSA.

9. Incubate the bead and antibody mix for 1 h at room temperature with gentle agitation and then move to 4°C with gentle agitation until needed.
10. Repeat the wash detailed in steps 21-24 to remove excess antibody.
11. Finally resuspend the beads in the original volume of 10 µL per organ to be collected (i.e., resuspend the beads for five organs in 50 µL of PBS/0.1% BSA).

### D. Tissue Digestion

1. Use petri dishes for organ preparations and ensure dissection tools are thoroughly cleaned between organ preparations to prevent cross contamination.
2. Use a new petri for each organ group and perform all digestion preparations in a tissue culture hood to keep cultures are sterile as possible. For digestion of lung **(i)**, liver **(ii)**, bone **(iii)** or brain **(iv)** see the relevant section below.

#### i) Lung Digestion

a) Place the lungs into a petri dish and wash once with cold PBS.
b) Remove any excess connective tissue including the trachea and bronchi using scissors and place this in a waste dish before washing the remaining lung tissue once more with cold PBS.
c) Mince the lung tissue with scissors first followed by a scalpel in a dry petri dish until tissue pieces are approximately 1 $mm^3$. Transfer the minced tissue to a 15 mL conical tube.
d) Add 10 mL PBS to the minced lungs with collagenase A to a final concentration of 2 mg/mL and parafilm the lid.
e) Digest the tissue at 37°C with gentle rotation for 2 h (Note: do not allow digestion to exceed 2 h).
f) Note: Adjust volume of digestion solution to number and volume of organs i.e., five lungs can be digested in 10 mL.

#### ii) Liver Digestion

a) Place the livers into a petri dish and wash once with cold PBS.
b) Check that the gall bladder has been removed, then gently separate the liver lobes and mince with scissors. Transfer the minced tissue to a 50 mL conical tube.
c) Add 15 mL PBS to the minced liver with collagenase A to a final concentration of 2 mg/mL and parafilm the lid.
d) Note: Adjust volume of digestion solution to number and volume of organs i.e., five livers can be digested in 15 mL.
e) Digest the tissue at 37°C with gentle rotation for 2 h (Note: do not allow digestion to exceed 2 h).

iii) **Bone Digestion**

a) Clean the femurs by removing any remaining muscles or tendons. Place the bones in sterile gauze to carefully rub away any attached soft tissue.
b) Place the bones in a mortar with 1 mL PBS and gently crush with a pestle to release the bone marrow.
c) Collect the PBS containing bone marrow and place in a clean 15 mL conical tube.
d) Add 1 mL of fresh PBS to the bone fragments remaining in the mortar and repeat crushing step with increased force. Collect the PBS containing bone marrow into the same collection conical tube. Note: PBS containing bone marrow will become lighter in colour with each wash/crush.
e) Repeat step 45 one more time with increased force until only small bone fragments remain.
f) Move the PBS and bone fragments from the mortar into the collection conical tube.
g) Make up the volume with PBS to the desired digestion volume i.e., five pairs of femurs in 10 mL. Add collagenase A to a final concentration of 2 mg/mL to the collection tube and parafilm the lid.
h) Digest the tissue at 37°C with gentle rotation for 1 h (Note: do not allow digestion to exceed 1 h).

iv) **Brain Digestion**

a) Place the brains in a clean, dry petri dish and use a scalpel to remove the brain stem, cerebellum, and olfactory bulb.
b) Roll the remaining cortical tissue on sterile blotting paper to remove the meninges.
c) Mince the remaining tissue with a scalpel until no large pieces remain. Transfer the tissue to a 15 mL conical tube and add 10 mL of pre-warmed DMEM-LG.
d) Break up the tissue further using a 10 mL serological pipette by gently mixing up and down.
e) Add collagenase A to a final concentration of 2 mg/mL and DNase1 to a final concentration of 10 μg/mL and parafilm the lid.
f) Digest the tissue at 37°C with gentle rotation for 1 h (Note: do not allow digestion to exceed 1 h).

### E. Isolation of pericytes from lung/liver/bone

1. Collect the digested tissue from the 37°C digestion incubator.
2. Filter the digest through a 70 μm nylon mesh filter into a new conical tube.
3. Centrifuge the flow through at 260 x g for 5 min. Note: This is a good time to perform the bead washing step outlined in steps 21-28.
4. Aspirate the supernatant from the digest carefully, the pellet will be loose.
5. Resuspend the cell pellet in 10 mL PBS to wash.

6. Centrifuge the washed cell suspension at 260 x g for 5 min.
7. Aspirate the supernatant carefully and resuspend the in PBS/0.1% BSA and move to a clean conical tube. Note: Volume of PSB/0.1% BSA depends on number of organs i.e., 4 mL for five organs.
8. Add the washed pre-coated beads to the cleaned tissue digest e.g., for five lungs and three livers the washed beads are resuspended in 80 µL of PBS/0.1% BSA so that 50 µL of bead suspension will be added to the cleaned lung digest and 30 µL will be added to the cleaned liver digest.
9. Incubate the tissue digest and bead suspension for 1 h at 4°C with gentle agitation.

**F. Isolation of pericytes from brain**

1. Collect the digested tissue from the 37°C digestion incubator.
2. Centrifuge the digest at 280 x g for 5 min.
3. Discard the supernatant and resuspend the pellet in 10 mL of 20% BSA-DMEM.
4. Mix the suspension well by passing up and down through a 10 mL serological pipette 20 times. Note: pipette slowly to reduce bubbles.
5. Centrifuge the suspension at 1000 x g for 10 min. The suspension will separate into 3 layers: myelin, supernatant, cell pellet (top to bottom).
6. Remove the milky myelin layer at the top, taking care to remove the myelin from the sides of the conical tube as well. Discard the remaining supernatant.
7. Resuspend the pellet in 2 mL of DMEM-LG and filter it through a 70 µm nylon mesh strainer into a new conical tube. Wash the strainer with 1 mL of DMEM.
8. Add 13 mL of DMEM-LG to the old conical tube, collect and pass through the same strainer into the new tube.
9. Centrifuge the suspension at 280 x g for 5 min.
10. Discard the suspension and resuspend the pellet in 2 mL DMEM-LG and transfer to a new 15 mL conical tube.
11. Wash the old conical tube with 2 mL DMEM-LG and transfer this to the new conical tube.
12. Add Dispase II to a final concentration of 1 mg/mL and DNase1 to a final concentration of 10 µg/mL.
13. Incubate the suspension at 37°C for 30 min with gentle rotation.
14. Centrifuge the suspension at 280 x g for 5 min.
15. Discard the supernatant and resuspend the pellet in 5 mL DMEM-LG to wash.
16. Centrifuge the suspension at 280 x g for 5 min. Note: This is a good time to perform the bead washing step outlined in steps 21-28.
17. Discard the supernatant and resuspend the pellet in PBS/0.1% BSA and move to a new conical tube. Note: Volume of PBS/0.1% BSA depends on number of organs i.e., 4 mL for five organs.
18. Add the washed pre-coated beads to the cleaned tissue digest.
19. Incubate the suspension for 1 h at 4°C with gentle agitation.

### G. Plating the pericytes

1. During the 1 h incubation at 4°C, wash the collagen coated plates with PBS.
2. Aliquot the bead-cell suspension across 1.5 mL microcentrifuge tubes and place the tubes in the magnetic holder.
3. Wait 30-60 s to allow the beads to separate from the suspension then gently remove the suspension without disturbing the beads.
4. Without the magnet, add 1 mL of PBS/0.1% BSA to one microcentrifuge tube and resuspend the beads.
5. Use this 1 mL volume for subsequent resuspension across tubes, collecting all the beads into one microcentrifuge tube.
6. Return the microcentrifuge tube containing all the beads to the magnetic holder and perform two more washes with PBS/0.1% BSA.
7. After washing, resuspend the beads in 1 mL of complete growth medium, and do not return the microcentrifuge tube to the magnet.
8. Add 2 mL of complete growth medium per well of a 6-well plate that has been pre-coated with collagen and washed with PBS.
9. Split the 1 mL cell suspension between the wells of the 6-well plate so that one well represents one organ i.e., for five organs use five wells on the 6-well plate adding 200 µL of the 1 mL total cell suspension per well.
10. Move the plate to the incubator with the appropriate oxygen tension and do not disturb for 7 days to ensure adherence and reduce cell loss.

### H. Maintaining the pericyte cultures

1. 7 d after isolation, replace the medium on the pericyte cultures with fresh equilibrated complete medium and continue to change medium every 2-3 d until cells are ready to be used. Note: Pericyte colonies should be visible at this time and typically will be ready to use in experiments 10-14 d after isolation.
2. Trypsinise the cells when they become 70-80% confluent with 500 µL 0.05% trypsin-ethylenediaminetetraacetic acid (EDTA) for approximately 10 min at 37°C and neutralise the trypsin with 500 µL of complete growth medium. Note: With each trypsinisation beads will detach from the cells.
3. Collect and centrifuge the cell suspension at 260 x g for 5 min.
4. Discard the supernatant and resuspend the cells in growth arrest (GA) medium. Note: If the pericytes remain clumpy in the resuspension, filter through a 70 µm filter for accurate counting and seeding.
5. Count and seed pericytes onto new collagen coated plates in GA medium 24 h before experiments begin.
6. Return the pericytes to the appropriate incubator i.e., 10% $O_2$ for lungs, 5% $O_2$ for brain, bones, and liver.

### Supplementary Tables

#### Supplementary Table 1

| **Media 1 (M1)** | **Media 2 (M2)** | **Media 3 (M3)** |
|---|---|---|
| Pericyte Medium containing FBS and Pen/Strep (ScienCell; 1201) | Dulbecco's Modified Eagle's Medium – low glucose (Millipore Sigma; D6046) | Dulbecco's Modified Eagle's Medium – low glucose (Millipore Sigma; D6046) |
| 1X Endothelial cell growth supplement (R&D; 390599) | Ham's F12 Nutrient Mix (Thermo Fisher Scientific; 11765047) | Ham's F12 Nutrient Mix (Thermo Fisher Scientific; 11765047) |
| | HEPES (Thermo Fisher Scientific; 15630080) | HEPES (Thermo Fisher Scientific; 15630080) |
| | Sodium Pyruvate (Thermo Fisher Scientific; 11360070) | Sodium Pyruvate (Thermo Fisher Scientific; 11360070) |
| | MEM non essential amino acids (Thermo Fisher Scientific; 11140050) | MEM non essential amino acids (Thermo Fisher Scientific; 11140050) |
| | 20% FBS (GeminiBio; 900-208-500) | 20% FBS (GeminiBio; 900-208-500) |
| | 1% Pen/Strep (Thermo Fisher Scientific; 15070063) | 1% Pen/Strep (Thermo Fisher Scientific; 15070063) |
| | | 1X Endothelial cell growth supplement (R&D; 390599) |

**Supplementary Table 2. Primer and Probe sequences for RT-qPCR.**

| **Gene** | **Primer/Probe** | **Sequence 5' -> 3'** |
|---|---|---|
| ***Rplp0*** | Forward primer | CTCTCGCTTTCTGGAGGGTG |
| | Reverse primer | TCAGTCTCCACAGACAATGCC |
| ***Cspg4*** | Forward primer | CGACTGCTGACTACAGATGATG |
| | Reverse primer | TGAGGAACCACGAGACTAGAA |
| ***Acta2*** | Forward primer | CCCCTGAAGAGCATCGGACA |
| | Reverse primer | TGGCGGGGACATTGAAGGT |
| ***Pdgfrb*** | Forward primer | GAACGACCATGGCGATGAGA |
| | Reverse primer | GCATCGGATAAGCCTCGAACA |
| ***Myh11*** | Forward primer | TCCTTCCTGGGCATTCT |
| | Reverse primer | CCTCTCGCTGGTACTCTT |
| ***Klf4*** | Forward primer | CACCCACACTTGTGACTATG |
| | Reverse primer | GTGCCTGGTCAGTTCATC |
| ***Rplp0*** | Probe | 5TEX615/TCCAGAGGCACCATTGAAATTCTGAGT/3IAbRQSp |
| | Forward primer | CCTCCTTCTTCCAGGCTTTG |
| | Reverse primer | CCACCTTGTCTCCAGTCTTTATC |
| ***Klf4*** | Probe | 56-FAM/CTTTTCCAA/ZEN/CTCGCTAACCCACCA/3IABkFQ |
| | Forward primer | GACCTCCTGGACCTAGACTTTA |
| | Reverse primer | GAAGACGAGGATGAAGCTGAC |

**Supplementary Table 3. Materials required for the isolation of pericytes from murine tissues.**

| Component | Supplier | Catalogue Number | Comments |
|---|---|---|---|
| 0.05% trypsin-EDTA | Thermo Fisher Scientific | 25300062 | |
| 15 mL Conical Tube | Alkali Scientific Inc | CN5600-NEST | |
| 50 ml Conical Tubes | Alkali Scientific Inc | CN5602-NEST | |
| 6-well cell culture plates | Alkali Scientific Inc | TP9006 | |
| 70 mm nylon cell strainer | Fisher Scientific | 22-363-548 | |
| 96-well cell culture plate | Alkali Scientific Inc | TPN1096 | |
| 96-well clear bottom black cell culture plate | Fisher Scientific | 07-200-567 | Used for immunofluorescence |
| Acetic Acid | Millipore Sigma | A6283 | |
| Albumin, Bovine Serum | Millipore Sigma | 12657 | |
| BioTek Lionheart FX automated microscope | Agilent | LFXW-SN | LED cubes required |
| Centrifuge | Thermo Fisher Scientific | 75009505 | |
| $CO_2$ incubator | Thermo Fisher Scientific | 51033544 | Hypoxia cabinet is placed in this incubator. |
| $CO_2$ incubator | Thermo Fisher Scientific | 51033597 | Set at 10% $O_2$, 5% $CO_2$ for lung pericytes. |
| $CO_2$-$O_2$-N2 (5-5-90) Mix | Roberts Oxygen | | For equilibration of Coy Lab Hypoxia Chamber at 5% $O_2$ |
| Collagen from rat tail | Millipore Sigma | C7661 | |
| Collagenase A | Millipore Sigma | 11088793001 | |
| Dispase II | Millipore Sigma | 4942078001 | |
| DNase | Millipore Sigma | D4263 | |
| Dulbecco's Modified Eagle's Medium – low glucose | Millipore Sigma | D6046 | |
| Dynabeads Sheep Anti-Rat IgG | Thermo Fisher Scientific | 11035 | Supplied at $4x10^8$ Dynabeads/mL |

| | | | |
|---|---|---|---|
| Endothelial cell growth supplement | R&D | 390599 | |
| Fetal Bovine Serum | GeminiBio | 900-208-500 | Lot tested (Lot No. A120030) |
| Graefe Forceps | Roboz Surgical Store | RS-5135 | |
| Ham's F12 Nutrient Mix | Thermo Fisher Scientific | 11765047 | |
| Heparin | Millipore Sigma | H3149 | |
| HEPES (1M) | Thermo Fisher Scientific | 15630080 | |
| Hypoxia cabinet with $O_2$ controller | Coy Lab Products | | Hypoxia unit is placed inside a standard tissue culture incubator (51033544). |
| Magnetic Separation Rack, 1.5 mL tubes | EpiCypher | 10-0012 | |
| MEM non-essential amino acids (100x) | Thermo Fisher Scientific | 11140050 | |
| Micro dissecting scissors | Roboz Surgical Store | RS-5906SC | |
| Ng2 Monoclonal Antibody (546930) | Thermo Fisher Scientific | MA5-24247 | Resuspend at 0.5 µg/µL |
| Nitrogen | Roberts Oxygen | | For equilibration of 10% oxygen incubator |
| Nutating Mixer | Corning | 6720 | |
| PBS, pH7.4 | Thermo Fisher Scientific | 10010023 | |
| Penicillin-Streptomycin | Thermo Fisher Scientific | 15070063 | |
| Petri Dish | Alkali Scientific Inc | TD0100 | |
| Sodium Pyruvate (100 mM) | Thermo Fisher Scientific | 11360070 | |
| Sorvall X4 Pro Centrifuge | Thermo Fisher Scientific | 75009505 | |
| Zeiss Primovert inverted cell culture microscope with Axiocam 208 | Zeiss | 415510-1101-000 | Camera attached |

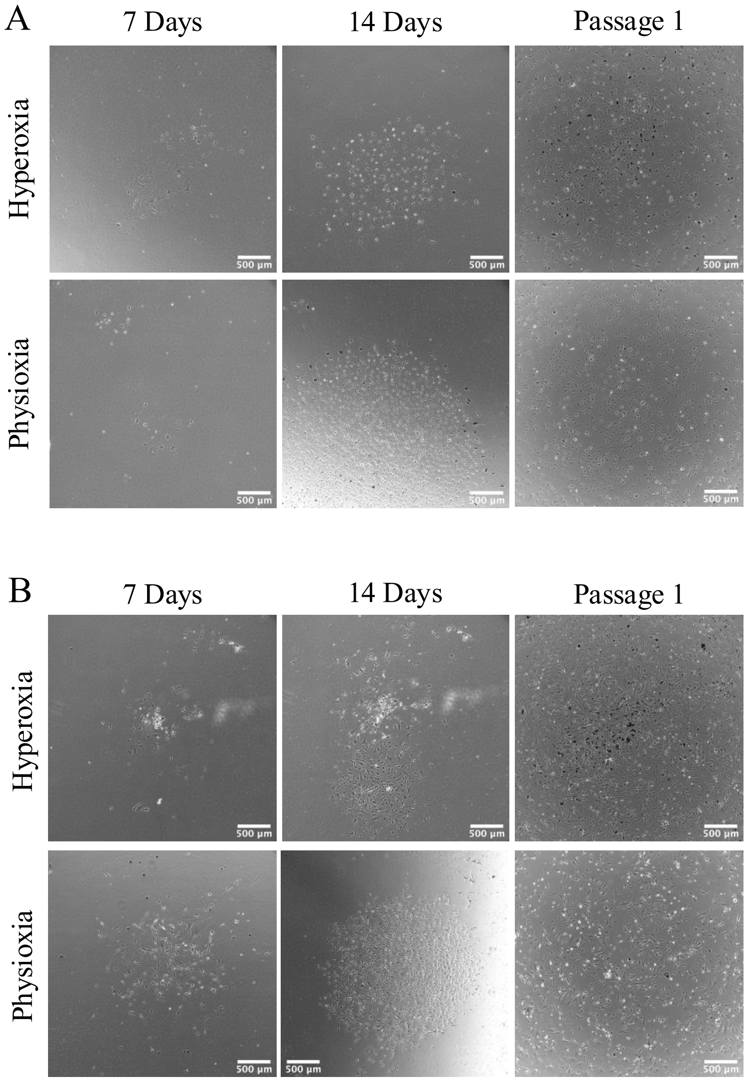


**Supplemental Figure 1. Physioxia supports primary lung pericyte expansion *ex vivo*.** Representative phase contrast images of primary lung pericytes expanded in medium composition M1 **(A)** or M2 **(B)** and maintained in either hyperoxia (21% $O_2$) or lung physioxia (10% $O_2$). Key time points during the expansion of primary pericytes are displayed. Scale bar = 500 µm.

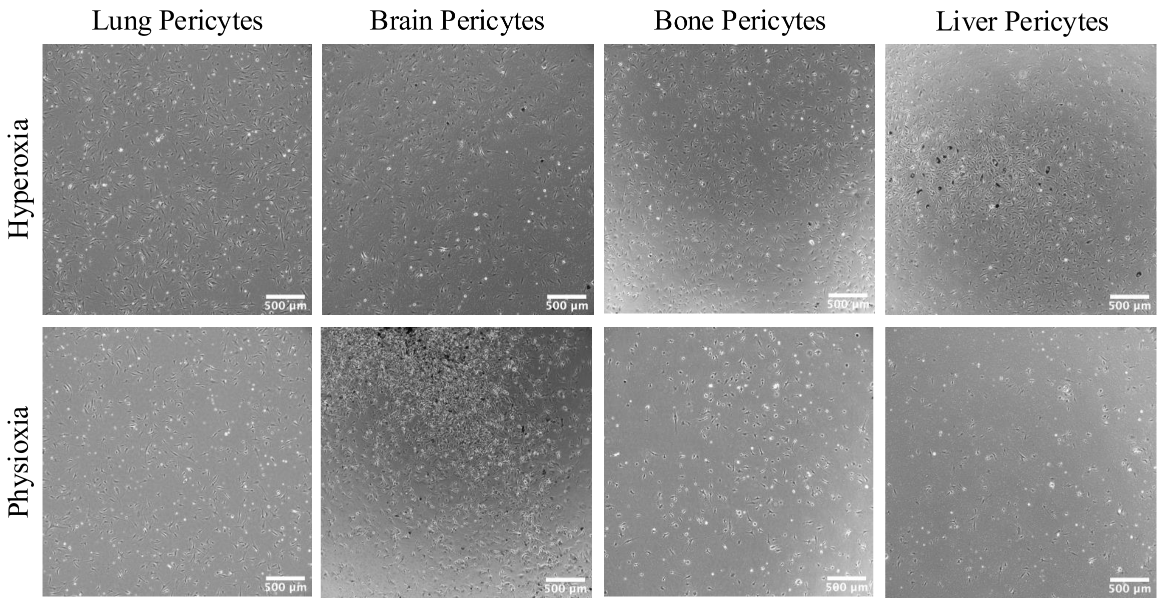


**Supplemental Figure 2. Physioxia retains characteristic pericyte morphology.** Representative phase contrast images of primary lung, brain, bone, and liver pericytes expanded in medium composition M3 and maintained in either hyperoxia (21% $O_2$) or physioxia (10% $O_2$ for lung; 5% $O_2$ for brain, bone, and liver). Scale bar = 500 μm.

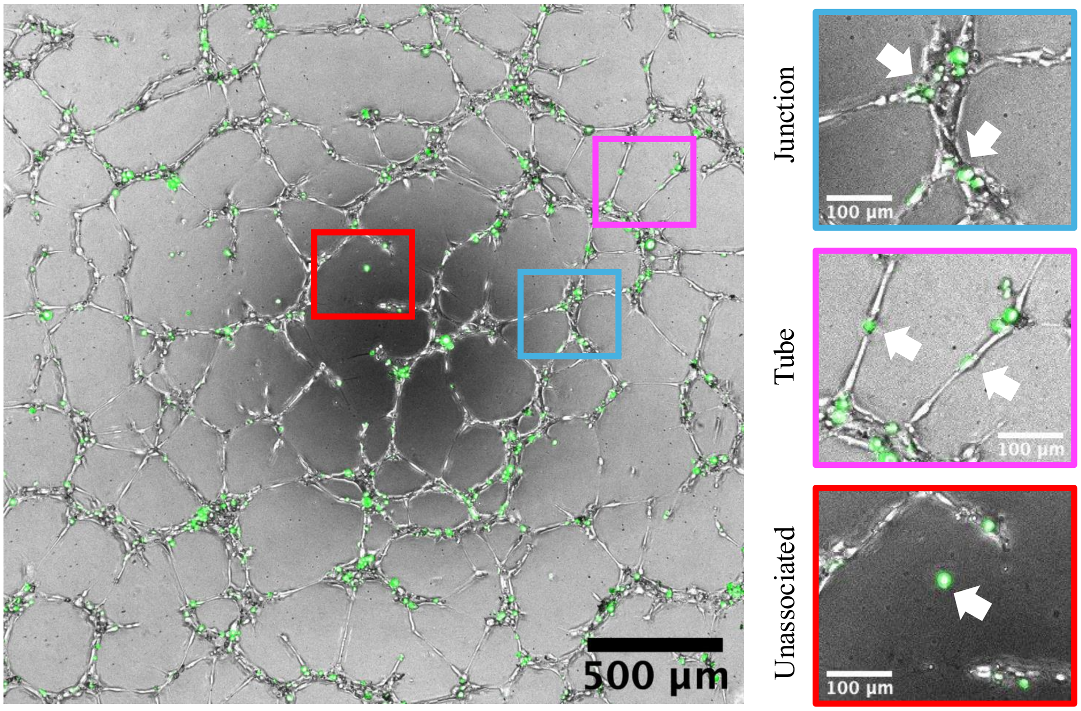


**Supplemental Figure 3. Pericyte localization on endothelial tubes.** Example image of hUVECs (unlabeled) co-cultured with zsGreen expressing brain pericytes at a ratio of 1:5 (PC:EC) in Matrigel after 4 h. Representative images of zsGreen expressing pericytes associating with endothelial junctions (blue), tubes (pink) or unassociated (red).

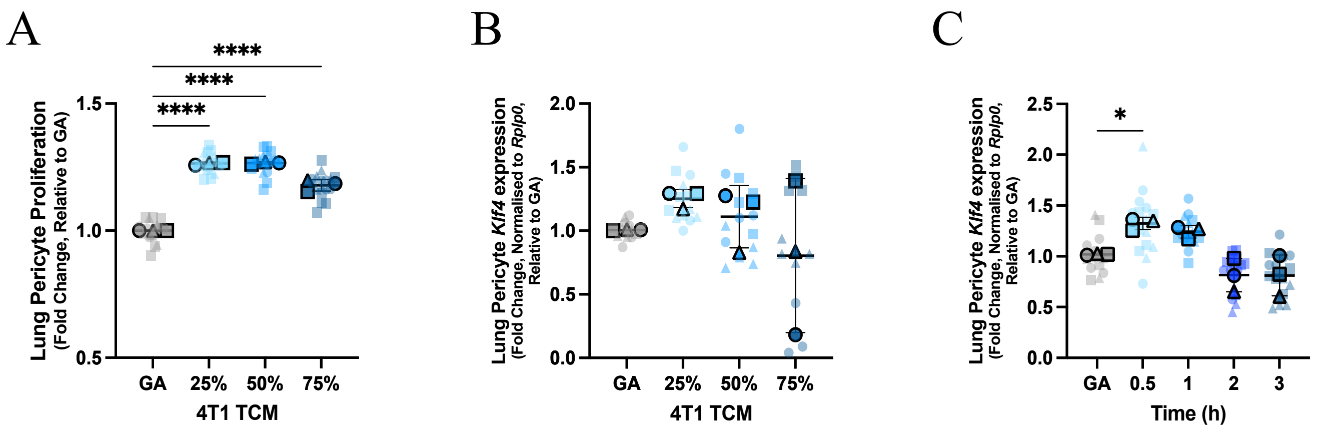


**Supplemental Figure 4. Defining lung pericyte activation conditions.** Lung pericytes were treated with various dilutions of 4T1 TCM to assess proliferation **(A)** at 72 h after treatment and *Klf4* expression **(B)** at 30 min after treatment. An activation time course **(C)** was performed to determine the peak *Klf4* expression time point after stimulation. Statistical significance was assessed by one-way ANOVA with Dunnett's multiple comparison test (*p<0.05, ****p<0.0001). Data are displayed as Superplots (Large symbols: means of experimental replicates; Small symbols: technical replicates for each experimental replicate; Lines: mean ±SD of replicate means).

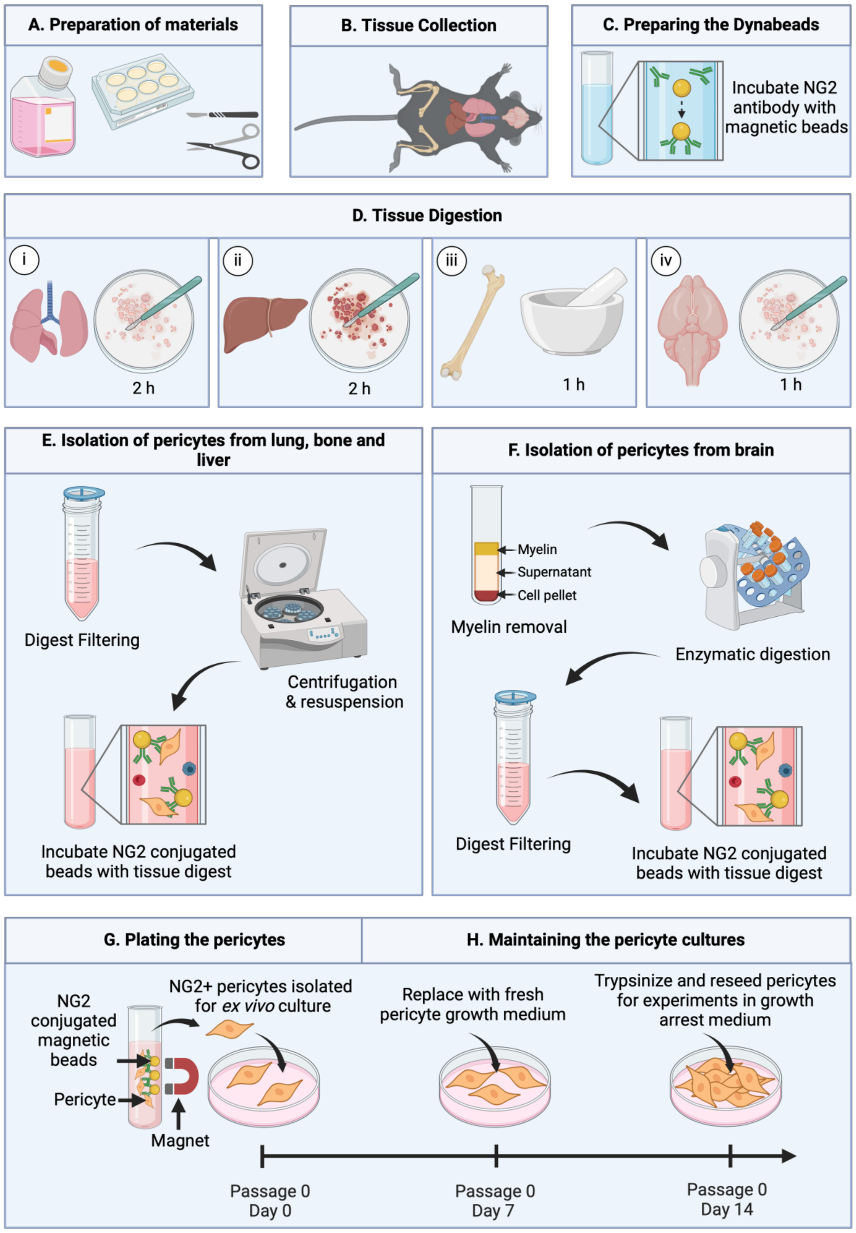


**Supplemental Figure 5. Pericyte isolation protocol. (A)** Prior to tissue collection, dissection tools are sterilized, pericyte medium is prepared and cell culture plates are coated with collagen. **(B)** Organs of interest are collected from 12 week old C57BL/6 female mice following euthanization by isoflurane and cervical dislocation. **(C)** Antibody conjugated beads are prepared by incubating magnetic Dynabeads with NG2 antibody. **(D)** Tissues undergo organ specific digestion protocols. **(E)** Tissue digests from lung, bone, and liver are filtered, centrifuged and incubated with NG2-conjugated beads which will bind to NG2-expressing pericytes. **(F)** Brain tissue digests undergo myelin removal and an additional digestion step before being filtered, centrifuged, and incubated with NG2-conjugated beads. **(G)** The bead bound NG2-expressing cells are isolated and washed using a magnet and plated for expansion under optimized pericyte culturing conditions **(H)**. Steps **(A-H)** correspond to the appropriate pericyte isolation protocol sections in supplemental materials. Figure made in Biorender.com.
